# Shared dynamic functional connectivity across schizophrenia, bipolar disorder and major depressive disorder

**DOI:** 10.1101/670562

**Authors:** Chao Li, Ke Xu, Mengshi Dong, Yange Wei, Jia Duan, Shaoqiang Han, Ruiqi Feng, Luheng Zhang, Pengfei Zhao, Yifan Chen, Xiaowei Jiang, Shengnan Wei, Zhiyang Yin, Yifan Zhang, Huafu Chen, Yanqing Tang, Fei Wang

## Abstract

Dynamic functional connectivity (DFC) analysis can capture time-varying properties of connectivity and may provide further information about transdiagnostic psychopathology across major psychiatric disorders. In this study, we used resting state functional MRI and a sliding-window method to study DFC in 150 schizophrenia (SZ), 100 bipolar disorder(BD), 150 major depressive disorder (MDD), and 210 healthy controls (HC). DFC were clustered into two functional connectivity states. Significant 4-group differences in DFC were found only in state 2. Post hoc analyses showed that transdiagnostic dysconnectivity among there disorders featured decreased connectivity within visual, somatomotor, salience and frontoparietal networks. Our results suggest that decreased connectivity within both lower-order (visual and somatomotor) and higher-order (salience and frontoparietal) networks may serve as transdiagnostic marker of these disorders, and that these dysconnectivity is state-dependent. Targeting these dysconnectivity may improve assessment and treatment for patients that having more than one of these disorders at the same time.

## Introduction

The traditional view of psychiatry holds that major psychiatric disorders (e.g., SZ, BD and MDD) are separate diagnostic categories with distinct etiologies and clinical presentations. However, existing diagnostic categories are not clearly associated with distinct neurobiological abnormalities [1; 2], which may hinder the search for biomarkers in psychiatry [3]. Major psychiatric disorders have common abnormalities in many characteristics, including genetic risk and etiology [4; 5], neural alterations [6-8], and clinical symptoms [9-11]. In addition, comorbidity among psychiatric disorders is very common, with 22% of patients having 2 diagnoses and 23% having 3 or more diagnoses [12]. Taken together, the abovementioned findings suggest that there are no clear-cut boundaries between different mental disorders. In contrast, each distinct psychiatric disorder is hypothesized to have broadly shared etiologies and mechanisms relative to the other psychiatric disorders [13; 14]. Transdiagnostic studies are necessary because they focus on fundamental processes underlying multiple psychiatric disorders, help to explain comorbidity among disorders, and may lead to improved assessment and treatment of disorders [15-17].

An important application of transdiagnostic models of psychopathology is to uncover shared (common) neurobiological abnormalities across multiple psychiatric disorders. The vast majority of previous studies, however, have merely compared patients with one specific group of psychiatric disorders to healthy controls (HC). Of the small number of transdiagnostic studies, many were meta-analyses that were based on individual studies using different methodologies [6; 8; 18; 19]. Recently, a small but rapidly growing number of original transdiagnostic studies have been conducted to directly investigate the shared abnormalities in brain structure and functional connectivity across multiple mental disorders [20-25]. For example, a previous study reported that disruptions within the frontoparietal network may be a shared feature across both SZ and affective psychosis [20]; this finding was validated and extended by a recent study [25]. In addition, a recent study found that the risk of common mental illnesses mapped onto hyperconnectivity between the visual association cortex and both the frontoparietal and default-mode networks [23].

Despite the contribution of advancing transdiagnostic research with regard to studies using functional connectivity, these studies assumed that the functional properties of the brain during the entire fMRI scan were static rather than dynamic. In fact, interactions among large-scale brain networks are highly dynamic, and time-averaged or static connectivity provides limited information about the functional organization of neural circuits [26; 27]. Therefore, using time-varying or dynamic methods to investigate shared patterns of dysfunction in large-scale functional connectivity networks across major psychiatric disorders may provide further information about their psychopathology. Along these lines, a few transdiagnostic studies have used DFC to investigate the dynamic functional architecture of brain networks in healthy young adults or the neurobiological abnormalities associated with psychiatric disorders [28-32]. For example, a previous study showed that DFC can reveal connectivity abnormalities that are not observed in static functional connectivity, suggesting that DFC is more sensitive than static functional connectivity [31]. Recently, another transdiagnostic study demonstrated that DFC was quite reliable within participants (within and across visits) and could act as a fingerprint, identifying specific individuals from within a larger group [28]. Although these studies that used DFC were very important, all of them had relatively small sample sizes (especially the patient groups) or studied only two diseases (i.e., SZ and BD or MDD and BD).

Here, we employed a widely used sliding-window approach [26; 27; 33; 34] to characterize DFC networks among patients with SZ (N = 150), BD (N = 100), or MDD (N = 150) and HC (N = 210). We aim to investigate the differences in dynamic connectivity between HCs and patients with SZ, BD or MDD. We hypothesized that the three disorders would share dysconnectivity in many brain networks.

## Results

### Two dynamic functional connectivity states

We used a sliding-window approach to construct the dynamic connectivity network. Then, we identified two patterns of dynamic connectivity network states (state 1 and state 2) using the k-means clustering method. State 1 was the less frequent of the two (35%) and featured stronger positive and negative connectivity. State 2 was more frequent (65%) and featured moderate positive and negative connectivity. The two states (represented by the centroids of the clusters) are shown in Figure 1.

**Fig. 1.**
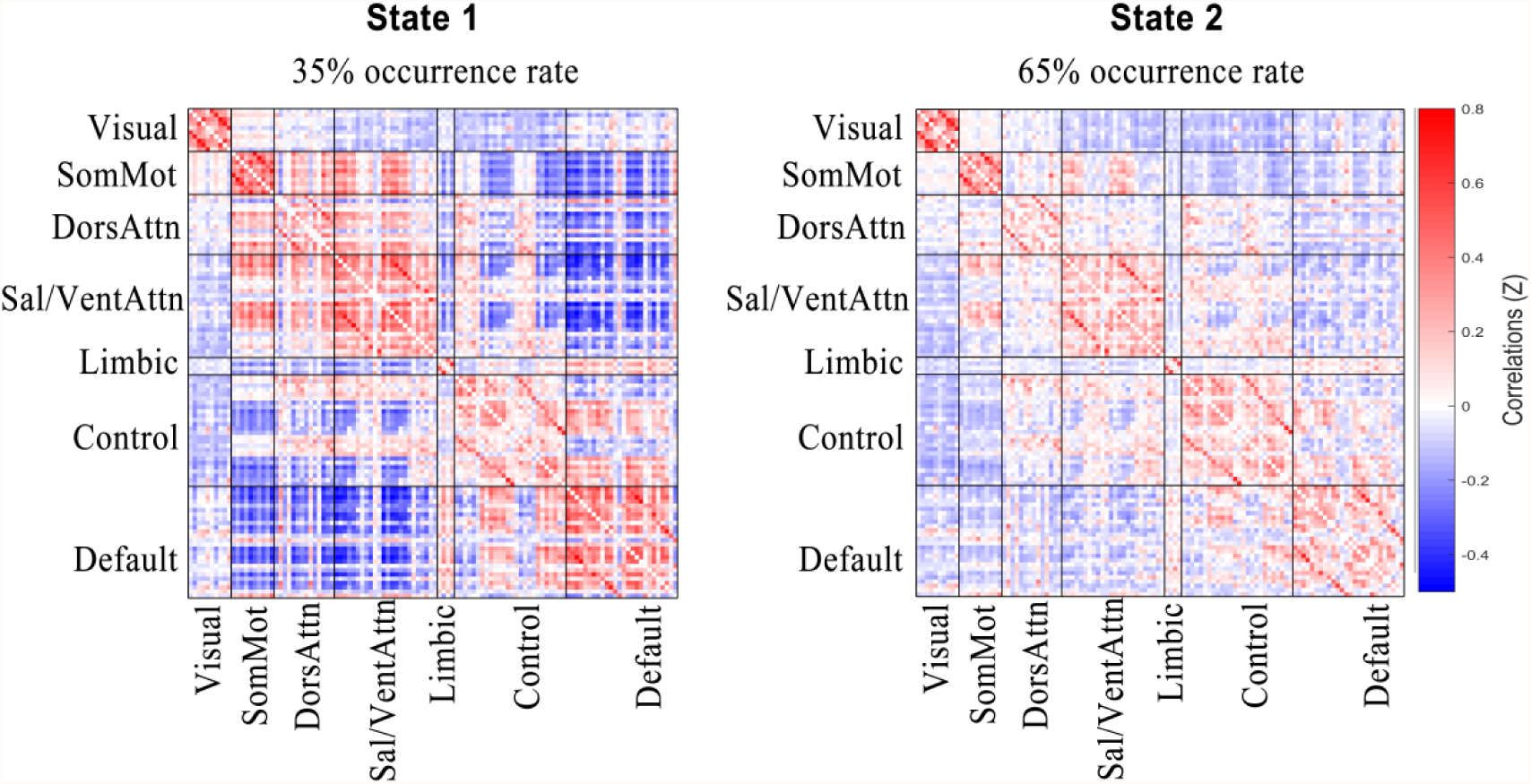
Cluster medians for each state. The total number of occurrences and the percentage of total occurrences are listed above each cluster median. The color bar represents the z value of dynamic functional connectivity. SomMot = somatomotor; DorsAttn = dorsal attention; Sal/VentAttn = salience/ventral attention; Control = frontoparietal control; Default = default mode.

### Differences in dynamic functional connectivity

Significant 4-group differences occurred only in state 2 (Figure 2.A; analysis of covariance [ANCOVA], FDR q < 0.05). Post hoc analyses in the state 2 identified significant dysconnectivity between patient group (i.e., SZ, BD and MDD) and HC (Figure 2.B-D; two-sample t-test, FDR q < 0.05). In general, SZ was more serious in both extent and range than those in BD and MDD. Figure S1 shows the group differences without statistical thresholding. To illustrate the pattern of differences at the level of the brain network, we present the average t values of the dysconnectivity within and between networks (Figure 3).

**Fig. 2.**
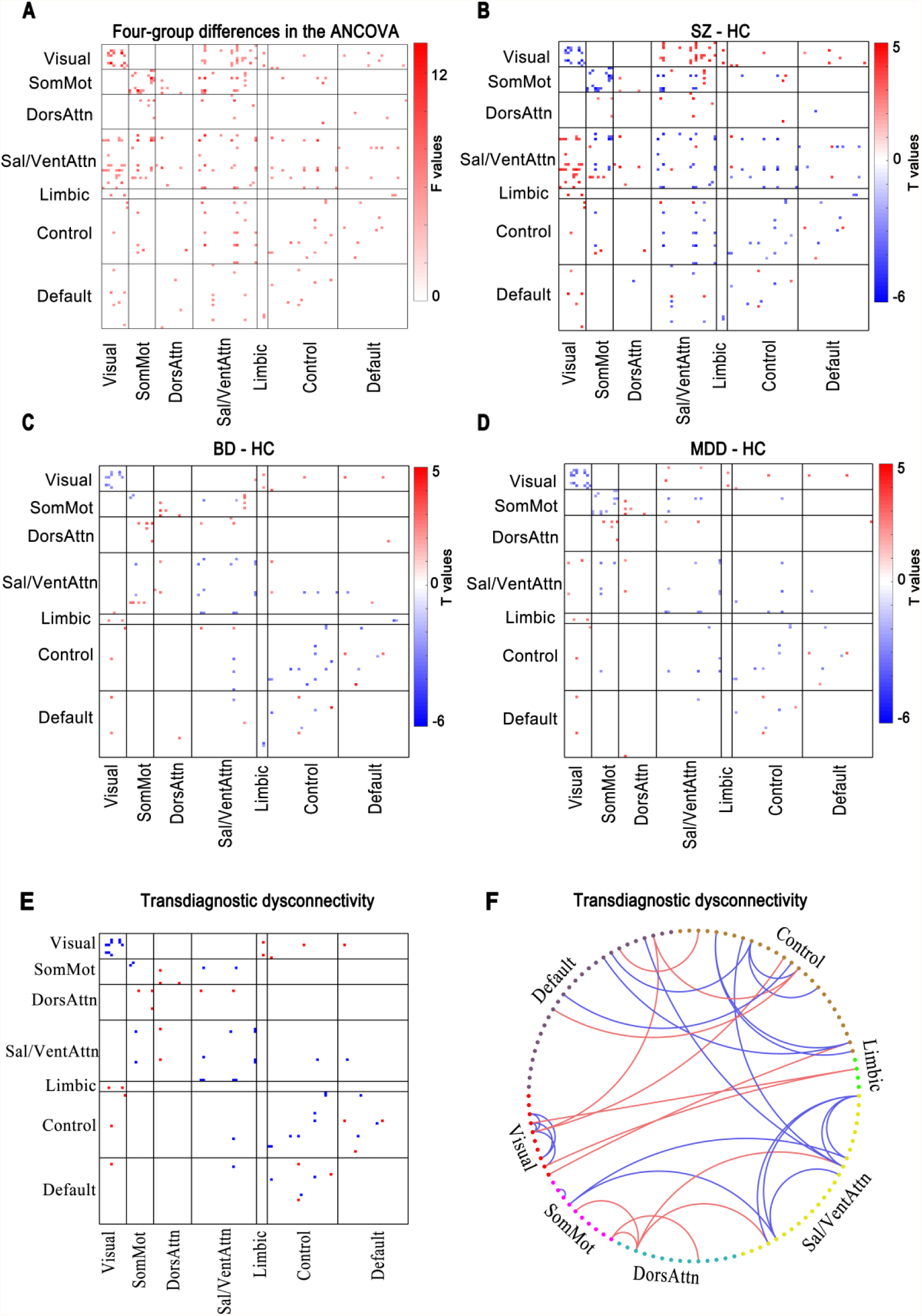
Significant differences in dynamic functional connectivity in state 2. (A) Four-group differences among schizophrenia, bipolar disorder, major depressive disorder and healthy controls (ANCOVA, FDR-corrected q < 0.05). (B-D) Group differences between patients and healthy controls (two-sample t-test, FDR-corrected q < 0.05). (E-F) Transdiagnostic dysconnectivity across these 3 disorders. The red dots or lines indicate increased connectivity, and the blue dots or lines indicate decreased connectivity. HC = healthy controls; SZ = schizophrenia; BD = bipolar disorder; MDD = major depressive disorder; SomMot = somatomotor; DorsAttn = dorsal attention; Sal/VentAttn = salience/ventral attention; Control = frontoparietal control; Default = default mode.

**Fig. 3.**
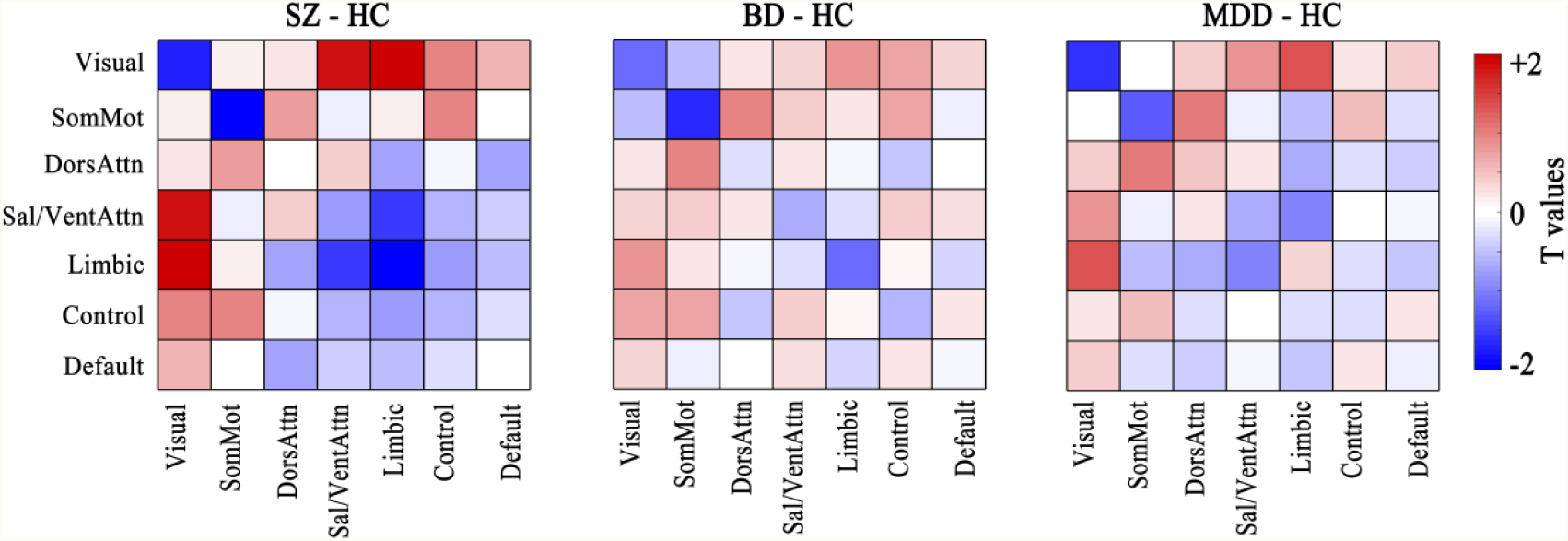
Average t-values of functional connectivity within and between networks. HC = healthy controls; SZ = schizophrenia; BD = bipolar disorder; MDD = major depressive disorder; SomMot = somatomotor; DorsAttn = dorsal attention; Sal/VentAttn = salience/ventral attention; Control = frontoparietal control; Default = default mode.

Based on the post hoc analyses, we further identified significantly dysconnectivity common to the SZ, BD, and MDD groups (Figure 2.E-F). Interestingly, the patterns of dysconnectivity were consistent across the three psychiatric disorders, i.e., they were either higher or lower than that of HC. The shared dysconnectivity within networks presented a consistent pattern of decreased connectivity, while the shared dysconnectivity between networks presented a mixed pattern of increased and decreased connectivity. Specifically, patients shared decreased connectivity within the visual, somatomotor, salience and frontoparietal control networks. Several patterns of dysconnectivity between networks were also common to these disorders. Specifically, connectivity was increased between the visual network and the limbic, frontoparietal and default-mode networks; between the dorsal attention network and the salience and somatomotor network; and between three pairs of regions in the frontoparietal and default-mode networks, while decreased connectivity between salience and frontoparietal control, default-mode and somatomotor networks, as well as three pairs of connectivity between frontoparietal and default-mode networks.

## Discussion

The present study was the first to examine alterations of DFC in SZ, BD and MDD with a relatively large sample size at a single site. We found that all participants experienced two distinct functional connectivity states during the resting state scanning: state 1, a less frequent, ‘extreme’ state characterized by stronger positive and negative connectivity, and state 2, a more frequent, moderate state with weaker connectivity. However, the group differences in DFC were expressed only in state 2. Post hoc analyses identified that shared patterns of dysconnectivity were marked by consistently decreased connectivity within most networks (visual, somatomotor, salience and frontoparietal networks), and SZ was the most obvious in extent. These findings suggest that decreased connectivity within both lower-order (visual and somatomotor) and higher-order (salience and frontoparietal) networks may serve as transdiagnostic marker of SZ, BD and MDD, and that these dysconnectivity is state-dependent. This study shed new light on the current transdiagnostic knowledge for these disorders. Targeting these transdiagnostic dysconnectivity may potentially improve assessment and treatment for psychiatric patients that having more than one of these disorders at the same time.

In the present study, there was no difference between the psychiatric patients and HC in state 1. Dysfunctional connectivity in the patients was manifested only in state 2. Interestingly, all shared dysconnectivity within networks were consistently decreased. Specifically, we found decreased connectivity within the visual, somatomotor, salience and frontoparietal networks across these psychiatric disorders. The finding of dysconnectivity in the frontoparietal network was consistent with previous studies showing transdiagnostic disruptions in this network across multiple psychiatric disorders [25; 35] as well as in SZ patients [36-39], BD patients [40; 41] and MDD patients [42-44]. The frontoparietal network is the core hub for cognitive control, adaptive implementation of task demands and goal-directed behavior [45-48]. In addition to the frontoparietal network, we also found decreased connectivity in the salience network, another network that is important for executive function. These findings also matched those of earlier studies showing abnormalities in the salience network [6; 49]. Reduced intra-network integration (or intra-network modularity [50]) in the cognitive networks may be responsible for cognitive dysfunction, one of the most prominent transdiagnostic characteristics of psychiatric disorders [8; 16].

Although abnormalities in higher-order brain networks such as the frontoparietal network were the dominant findings in previous studies, a recent study [51] suggests that the focus of psychiatric neuroscience should be expanded beyond these networks to some lower-order networks, such as the somatomotor and visual networks. The present findings of dysconnectivity within the somatomotor and visual networks corroborated earlier studies showing abnormalities in these networks in psychiatric disorders [22; 23; 52]. Together, these findings suggested that intra-network integration was decreased in patients with psychiatric disorders not only within higher-order brain networks but also within lower-order brain networks. Future research should pay additional attention to lower-order networks.

In contrast, the inter-network connectivity of patients compared to HC was not consistently decreased, but increased for some connections and decreased for others. Increased connectivity was mainly driven by connectivity between the visual network and the limbic, frontoparietal and default-mode networks. Although dysconnectivity between the visual network and others was not often considered primary to psychopathological dysfunction in early studies, this finding was in line with that of a recent study [23] showing that hyperconnectivity between the visual association cortex and the frontoparietal and default-mode networks was correlated with increased p-factor scores (a single general transdiagnostic factor associated with risk for all common forms of mental illness). Considering that the patient groups in the present study were all psychiatric patients, our findings support and expand on that recent study, suggesting that there is dysconnectivity between the visual network and the frontoparietal and default-mode networks not only in those who are at a high risk for psychiatric disorders but also in those who already suffer from psychiatric disorders. This shared feature represents the possibility of a trait common to multiple psychiatric disorders.

Decreased connectivity was mainly driven by dysconnectivity between the salience network and the frontoparietal, default-mode and somatomotor networks. The salience, frontoparietal and default-mode networks are the three most important high-order cognitive networks. Many studies have indicated that the salience network plays a role in switching between the frontoparietal and default-mode networks to improve performance of cognitively demanding tasks [53; 54]. Abnormal communication between these networks may be one of the factors underlying cognitive impairment in psychiatric disorders. Our findings are consistent with a recent review [8].

Strikingly, the shared patterns of dysconnectivity across these disorders were all in the same direction, i.e., they were either higher or lower than those of HC. Although speculative, this finding may support the idea that SZ, BD and MDD may lie along a transdiagnostic continuum of major endogenous psychoses [55].

Our study has several limitations. The first limitation is the choice of window size for sliding-window analysis. In this study, we used an empirically validated fixed sliding-window of 17 TRs (34s) as suggested by previous studies [28; 56] to maximize signal estimates while still capturing the properties of transient functional connectivity. Future work should evaluate DFC across a variety of window sizes. Second, we did not record respiratory and cardiac events and use them for denoising, which may have had an impact on our results. Finally, this is a cross-sectional research, longitudinal study is needed to better understand the transdiagnostic pathophysiological mechanisms for these psychiatric disorders.

In conclusion, we performed, to our knowledge, the first time-varying functional connectivity analyses in HC and SZ, BD and MDD patients in a single study with a relatively large sample size. Functional connectivity disruptions in psychosis were state specific and intermittent. Importantly, the patterns of shared dysconnectivity were marked by consistently decreased connectivity within both higher-order and lower-order brain networks, while there was a mixed pattern of increased and decreased connectivity between distributed networks. Our findings expand the current understanding of the transdiagnostic pathophysiological mechanisms for these 3 psychiatric disorders. Targeting these shared dysconnectivity may potentially improve assessment and treatment for psychiatric patients that having more than one of these disorders at the same time.

## Methods

### Participants

The study was approved by the Institutional Review Board of China Medical University. All participants provided written informed consent after receiving a detailed description of the study. Eight hundred and fifty-one individuals participated in this study, including 332 HC, 183 SCZ patients, 132 BD patients and 204 MDD patients. We finally included 610 participants (see the **Head motion control** section for details), including 210 HC, 150 SZ patients, 100 BD patients and 150 MDD patients. The demographics, clinical characteristics, cognitive function and head motion information of the included participants are summarized in Table 1. All participants with SZ, BD, and MDD were recruited from the inpatient and outpatient services at the Shenyang Mental Health Center and the Department of Psychiatry at the First Affiliated Hospital of China Medical University, Shenyang, China, between February 2009 and April 2018. HC were recruited from the local community by advertisement.

**Table 1.** Demographics, Clinical Characteristics, and Cognitive Function of Healthy Controls and Patients with Schizophrenia, Bipolar Disorder, and Major Depressive Disorder.

| Variable | HC<br>(n =210) | SZ<br>(n =150) | BD<br>(n =100) | MDD<br>(n =150) | F/$\chi^2$ values | P values |
| --- | --- | --- | --- | --- | --- | --- |
| Demographic characteristic |  |  |  |  |  |  |
| Age at scan (y) | 24.37 (5.74) | 23.67 (8.77) | 24.56 (5.95) | 25.53 (8.30) | 1.663 | 0.174 |
| Male | 86 (41%) | 59 (39%) | 36 (36%) | 43 (29%) | 6.269 | 0.099 |
| Clinical characteristic |  |  |  |  |  |  |
| Duration (mo) | N/A | 23.27 (34.97) | 36.61 (36.09) | 20.57 (28.10) | 7.745 | 0.001 |
| First episode (yes) | N/A | 96 (64%) | 48 (48%) | 122 (81%) | 30.599 | 2.267E-7 |
| Medication (yes) | N/A | 111 (74%) | 65 (65%) | 85 (57%) | 9.941 | 0.007 |
| Antidepressants | N/A | 12 (8%) | 30 (30%) | 58 (39%) | 39.395 | 2.788E-9 |
| Antipsychotics | N/A | 71 (47%) | 28 (28%) | 6 (4%) | 72.958 | 1.110E-16 |
| Mood stabilizers | N/A | 5 (3%) | 39 (39%) | 2 (1%) | 99.370 | 0.000 |
| Anxiolytics | N/A | 14 (9%) | 7 (7%) | 31 (21%) | 12.762 | 0.002 |
| Other drugs | N/A | 9 (6%) | 0 (0%) | 0 (0%) | N/A | N/A |
| HAMD-17 | (n = 194) | (n = 121) | (n = 94) | (n = 147) |  |  |
|  | 1.06 (1.71) | 6.82 (6.14) | 11.92 (9.78) | 19.18 (9.84) | 187.866 | 4.151E-84 |
| HAMA | (n = 194) | (n = 113) | (n = 94) | (n = 135) |  |  |
|  | 0.92 (2.15) | 5.99 (6.22) | 9.56 (9.33) | 15.99 (9.78) | 127.235 | 3.937E-62 |
| YMRS | (n = 192) | (n = 113) | (n = 97) | (n = 135) |  |  |
|  | 0.17 (0.66) | 1.93 (4.70) | 6.72 (9.71) | 1.27 (2.88) | 40.150 | 2.095E-23 |
| BPRS | (n = 162) | (n = 147) | (n = 86) | (n = 98) |  |  |
|  | 18.40 (1.24) | 33.82 (13.11) | 26.20 (9.45) | 27.18 (7.39) | 78.791 | 1.362E-41 |
| Cognitive function |  |  |  |  |  |  |
| WCST | (n = 173) | (n = 107) | (n = 87) | (n = 114) |  |  |
| Correct responses | 33.09 (10.43) | 19.79 (11.44) | 28.10 (11.30) | 26.11 (11.20) | 33.111 | 1.865E-19 |
| Categories completed | 4.46 (1.90) | 1.88 (1.93) | 3.57 (1.97) | 3.15 (1.97) | 39.934 | 4.874E-23 |
| Total errors | 14.95 (10.59) | 28.21 (11.44) | 19.54 (11.23) | 21.85 (11.24) | 32.645 | 3.308E-19 |
| Perseverative errors | 5.30 (6.46) | 11.36 (9.71) | 7.74 (8.50) | 8.79 (8.71) | 12.624 | 5.919E-8 |
| Nonperseverative errors | 9.58 (5.55) | 16.79 (8.15) | 12.09 (6.15) | 13.06 (6.50) | 27.310 | 2.598E-16 |
| Head motion parameters |  |  |  |  |  |  |
| Mean FD, mm | 0.10 (0.03) | 0.10 (0.04) | 0.11 (0.04) | 0.10 (0.03) | 1.923 | 0.125 |
| Percentage of excessive FD | 0.07 (0.07) | 0.06 (0.06) | 0.08 (0.08) | 0.06 (0.06) | 1.805 | 0.145 |
| Max translation, mm | 0.44 (0.46) | 0.53 (0.55) | 0.50 (0.47) | 0.50 (0.44) | 1.080 | 0.357 |
| Max rotation, degree | 0.38 (0.34) | 0.40 (0.38) | 0.46 (0.37) | 0.46 (0.46) | 1.919 | 0.125 |

Note: Data are presented as either number (%) or means (standard deviations).
HC, healthy controls; SZ, schizophrenia; BD, bipolar disorder; MDD, major depressive disorder; HAMD, Hamilton Depression Scale; HAMA, Hamilton Anxiety Scale; YMRS, Young Mania Rating Scale; BPRS, Brief Psychiatric Rating Scale; WCST, Wisconsin Card Sorting Test; FD, framewise displacement.
N/A, not available/not applicable.

The presence or absence of Axis I psychiatric diagnoses was determined by two trained psychiatrists using the Structured Clinical Interview for Diagnostic and Statistical Manual of Mental Disorders, Fourth Edition (DSM-IV) Axis I Disorders for participants 18 years and older, whereas the Schedule for Affective Disorders and Schizophrenia for School-Age Children–Present and Lifetime Version (K-SADS-PL) was used for participants younger than 18 years. All patients with SZ, BD, or MDD were required to meet the DSM-IV diagnostic criteria for their respective disorders and no other Axis I disorders. The HC did not have a current or lifetime history of any Axis I disorder or a history of psychotic, mood, or other Axis I disorders in first-degree relatives, as determined from a detailed family history. Potential participants were excluded if they had (1) lifetime substance/alcohol abuse or dependence, (2) the presence of a concomitant major medical disorder, (3) any MRI contraindications, (4) a history of head trauma with loss of consciousness ≥5 minutes or any neurological disorder, or (5) any abnormality identified by T1- or T2-weighted imaging. Symptoms and cognitive measures were assessed using the Brief Psychiatric Rating Scale (BPRS), the Hamilton Depression Rating Scale (HAMD), the Hamilton Anxiety Rating Scale (HAMA), the Young Mania Rating Scale (YMRS), and the Wisconsin Card Sorting Test (WCST).

### MRI acquisition

MRI data were acquired using a GE Signa HD 3.0-T scanner (General Electric, Milwaukee, WI) with a standard 8-channel head coil at the First Affiliated Hospital of China Medical University. Functional imaging was performed using a gradient-echo-planar imaging (EPI-GRE) sequence. The following parameters were used: repetition time = 2000 ms, echo time = 30 ms, flip angle = 90°, field of view = 240 mm × 240 mm, matrix = 64 × 64, slice thickness = 3 mm with no gap, number of slices = 35. The scan lasted 6 minutes and 40 seconds, resulting in 200 volumes. Participants were instructed to rest and relax with their eyes closed but to remain awake during the scanning.

### Data preprocessing

All images were preprocessed using SPM12 (www.fil.ion.ucl.ac.uk/spm/) and Data Processing & Analysis of Brain Imaging (DPABI) [57]. The volumes from the first 10 time points were discarded. The subsequent preprocessing steps included slice-timing correction and head motion correction. The corrected functional images were spatially normalized to the Montreal Neurological Institute space using the EPI template in SPM12, resampled to 3 mm × 3 mm × 3 mm isotropic voxels, and further smoothed via a Gaussian kernel with a 4-mm full width at half-maximum. Then, we performed linear detrending and temporal bandpass filtering (0.01–0.01 Hz) to reduce low-frequency drift and high-frequency noise. Next, several confounding covariates, including the Friston-24 head motion parameters, white matter, cerebrospinal fluid, and global signals, were regressed out of the blood oxygen level-dependent (BOLD) time series for all voxels.

### Head motion control

Because excessive head motion can significantly affect dynamic connectivity analysis [58; 59], we carried out head motion control and discarded participants with excessive head motion before dynamic connectivity analysis. We discarded participants if they had mean framewise displacement (FD) values > 0.2 mm, if the outliers accounted for > 30% of all volumes (190 volumes), or if head motion exceeded 3 mm or 3°. According to these criteria, we excluded 41 HC, 30 SZ, 18 BD and 25 MDD patients. To match the four groups by age and sex, we further excluded 81 HC patients (including 2 HC patients who lacked age and gender information), 3 SZ patients, 14 BD patients and 29 MDD patients. The mean FD, percentage of outliers and maximum head motion are presented in Table 1.

### Dynamic functional connectivity analysis

The average BOLD time series of 114 nodes within the 17-network functional atlas of Yeo et al. [60] were extracted. Dynamic connectivity was estimated from these time series with a widely used sliding-window approach in the software GIFT [61; 62]. The window was a rectangular window of 17× the repetition time (TR) convolved with a Gaussian of sigma 3×TR to obtain a tapered window, and it slid in steps of 1 TR. A previous study [56] suggested that a sliding-window width range of 30–60 s was appropriate for dynamic connectivity analyses. This previous study also revealed consistent state solution stability across varying sliding-window sizes of 33–63 s. Consequently, a width of 17×TR (i.e., 34 s) was chosen to maximize signal estimates while still capturing the properties of transient functional connectivity. As previous studies suggest that covariance estimation using shorter time series can be noisy, we estimated covariance from the regularized precision matrix (inverse covariance matrix) [63; 64]. Furthermore, we imposed a penalty on the L1 norm of the precision matrix to promote sparsity using the graphical least absolute shrinkage and selection operator (LASSO) method [65]. For each participant, the regularization parameter lambda was optimized by evaluating the log-likelihood of unseen data from the same subject in a cross-validation framework. For each participant, we obtained a total of 173 windows, each of which had (114×113)/2 = 6,441 unique functional connectivity measurements. Finally, Fisher r-to-z transformation was performed for all functional connectivity measurements.

### State clustering analysis

The k-means algorithm can identify sets of time-varying network configurations in different windows that have common features, grouping them into clusters that are more similar to each other than to configurations in other clusters. We used the Manhattan distance (*L*_1_ distance) as a similarity measure in clustering, as it has been demonstrated to be the most effective measure for high-dimensional data [66]. To reduce the computational demands and to diminish redundancy between windows, following a previous study [26], we first used the subject exemplars as a subset of windows with local maxima in functional connectivity variance to perform k-means clustering with varying numbers of clusters k (2–10). The optimal number of clusters k = 2 was determined based on the silhouette criterion [67], a cluster validity index that reflects how similar a point is to other points in its own cluster compared to points in other clusters. The resulting 2 cluster centroids were used as starting points to cluster all DFC data (610 subjects × 173 windows=105,530 matrices) into 2 clusters. The finally resulting cluster centroids were regarded as functional connectivity states at the group level. For each participant, each state was regarded as the median of that windowed functional connectivity that had the same cluster index (i.e., 1 and 2).

### Statistical analysis

Group effects on dynamic connectivity were examined using one-way ANCOVA, with age, sex and mean FD as covariates. Regarding post hoc analyses, two-sample t-tests were performed following significant group effects with ANCOVA, which were compared in a pairwise fashion with the HC group as the common comparison. We used the FDR to correct for multiple comparisons in both the ANCOVA and the post hoc analyses (q < 0.05). Shared dysconnectivity was defined as a situation in which all three patient groups had abnormal connectivity compared with the HC group.

## Code availability

All analysis code is available here: https://github.com/lichao312214129/lc_rsfmri_tools_matlab/tree/master/Workstation/code_workstation2018_dynamicFC

## Acknowledgements

This study was funded by the National Science Fund for Distinguished Young Scholars (81725005 to F.W.), the National Natural Science Foundation of China (81571311 to Y.T., 81571331 to F.W.), the National Key Research and Development Program (2018YFC1311604 to Y.T., 2016YFC1306900 to Y.T., 2016YFC0904300 to F.W.), the National High Tech Development Plan (863) (2015AA020513 to F.W.), the Liaoning Science and Technology Project (2015225018 to Y.T.), the Liaoning Education Foundation (Pandeng Scholar to F.W.), the Innovation Team Support Plan of Higher Education of Liaoning Province (LT2017007 to F.W.), and the Major Special Construction Plan of China Medical University (3110117059 to F.W.). This manuscript has been released as a preprint on bioRxiv [68].

## Notes

#### Summary of Updates

Revison 1: We have reprocess all our fmri data. This time we did not do the despiking. We used a rigorous method to control the head motion: subjects with greater head motion were excluded from this study (Please see Head motion control section of the revised manuscript). We discarded participants if they had mean framewise displacement (FD) values > 0.2 mm, if the outliers accounted for > 30% of all volumes (190 volumes), or if head motion exceeded 3 mm or 3 degree. According to these criteria, we excluded 41 healthy controls, 30 patients with schizophrenia, 18 patients with bipolar disorder and 25 patients with major depressive disorder. Besides, we treated the age, sex and mean FD as covariates, when we performed ANCOVA. Revison 2: This asymmetry is because we only extract the lower triangular matrix in the process of data analysis, and when we get results, we do not mirror the data of the lower triangular matrix into the upper triangular matrix. In this way, when sorting nodes and edges according to their brain network index, this asymmetry emerged. In the revised version, we mirror the lower triangular matrix into the upper triangular matrix (using the MATLAB command M = M + M'; M is the lower triangular matrix), which solves this bug. Revison 3: We reclustered the dynamic functional connectivity: "we used the Manhattan distance (L1 distance) as a similarity measure in clustering, as it has been demonstrated to be the most effective measure for high dimensional data. To reduce the computational demands and to diminish redundancy between windows, we first used the subject exemplars as a subset of windows with local maxima in functional connectivity variance to perform kmeans clustering with varying numbers of clusters k (from 2 to 10). The optimal number of clusters k = 2 was determined based on the silhouette criterion, a cluster validity index that reflects how similar a point is to other points in its own cluster compared to points in other clusters." Revison 4: According to this suggestion, we additionally showed group differences without statistical thresholding (Supplementary_material Figure S1), so that the interpretation is not completely driven by the choice of statistical threshold. Revison 5: We have presented the head motion information in Table 1. After discarded those participants with greater head motion, there was indeed no statistical difference between the four groups (ANOVA p < 0.05).

https://github.com/lichao312214129/lc_rsfmri_tools_matlab/tree/master/Workstation/code_workstation2018_dynamicFC

