## Supplementary material for "Shared dynamic functional connectivity across schizophrenia, bipolar disorder and major depressive disorder"

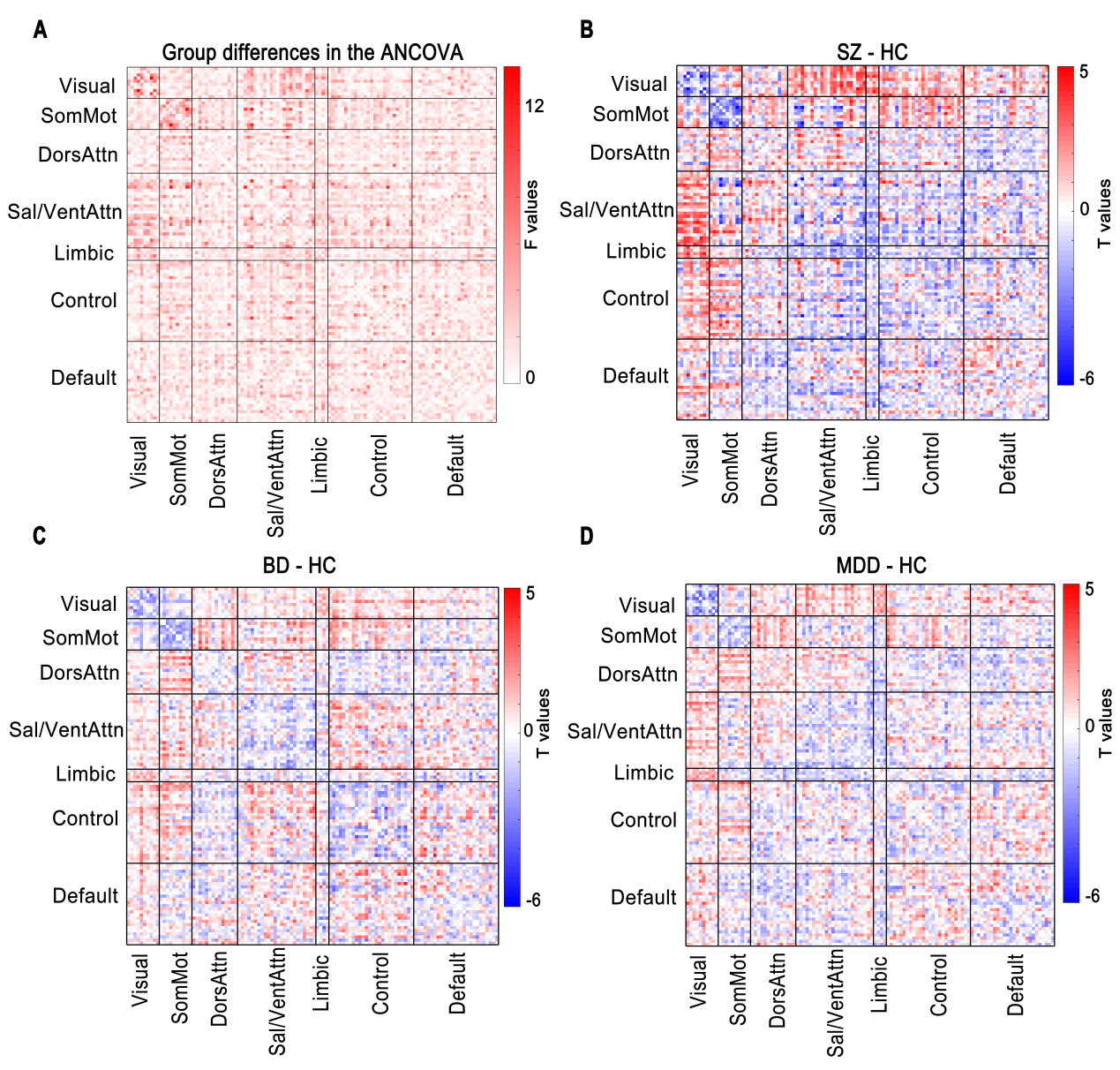

**Fig. S1 Four-group differences in dynamic functional connectivity in state 2 without statistical thresholding.** (A) Group differences among schizophrenia patients, bipolar disorder patients, major depressive disorder patients and healthy controls obtained using ANCOVA. (B-D) Group differences between patients and healthy controls. If the ANCOVA revealed significant group effects, the differences between groups were evaluated using a post hoc pairwise two-sample t-test. HC = healthy controls; SZ = schizophrenia; BD = bipolar disorder; MDD = major depressive disorder; SomMot = somatomotor; DorsAttn = dorsal attention; Sal/VentAttn = salience/ventral attention; Control = frontoparietal control; Default = default mode.
